# Positive selection inhibits plasmid coexistence in bacterial genomes

**DOI:** 10.1101/2020.09.29.318741

**Authors:** Laura Carrilero, Anastasia Kottara, David Guymer, Ellie Harrison, James P. J. Hall, Michael A. Brockhurst

**Affiliations:** Division of Evolution and Genomic Sciences, School of Biological Sciences, University of Manchester, Manchester, M13 9PT, UK; Department of Animal and Plant Sciences, University of Sheffield, Sheffield, S10 2TN, UK; Department of Biology, University of York, York, YO10 5DD, UK; Department of Evolution, Ecology and Behaviour, Institute of Integrative Biology, University of Liverpool, Liverpool, L69 7BZ, UK

## Abstract

Plasmids play an important role in bacterial evolution by transferring niche adaptive functional genes between lineages, thus driving genomic diversification. Bacterial genomes commonly contain multiple, coexisting plasmid replicons (i.e., plasmid coinfection), which could fuel adaptation by increasing the range of gene functions available to selection and allowing their recombination. However, plasmid coinfection is difficult to explain because the acquisition of plasmids typically incurs high fitness costs for the host cell. Here, we show that plasmid coinfection was stably maintained without positive selection for plasmid-encoded gene functions and was associated with compensatory evolution to reduce fitness costs. By contrast, with positive selection, plasmid coinfection was unstable despite compensatory evolution. Positive selection discriminated between differential fitness benefits of functionally redundant plasmid replicons, retaining only the more beneficial plasmid. These data suggest that while the efficiency of negative selection against plasmid fitness costs declines over time due to compensatory evolution, positive selection to maximise plasmid-derived fitness benefits remains efficient. Our findings help to explain the forces structuring bacterial genomes: Coexistence of multiple plasmids in a genome is likely to require either rare positive selection in nature or non-redundancy of accessory gene functions among coinfecting plasmids.

## Introduction

Plasmids play an important role in the evolution of bacterial genomes, promoting evolutionary divergence by transferring niche-adaptive accessory gene functions between lineages[1–3]. Bioinformatic analyses suggest that carriage by single bacterial host cells of multiple, coexisting plasmid replicons—i.e. plasmid coinfection—is more commonly observed than would be expected by chance[4]. Accumulation of multiple, coexisting plasmid replicons thus drives genome expansion and can lead to the evolution of multipartite genomes [5–7]. Moreover, this process could potentially fuel evolutionary innovation through reassortment of new combinations of accessory functions and the formation of new mobile genetic elements through plasmid-plasmid recombination[8, 9]. Nevertheless, the observed high rates of plasmid coinfection are surprising given that plasmid acquisition usually disrupts normal cellular function and is often associated with large fitness costs for the host cell[10–14].

One explanation for abundant plasmid coinfection is that the fitness costs of acquiring multiple plasmids could be less than additive. Positive epistasis between plasmid costs could permit the accumulation of multiple plasmids by reducing the cost for plasmid-bearers of acquiring additional plasmids[4, 15, 16], however, positive epistatic interactions among plasmid costs are not universal[4]. Moreover, as we will show in this study, the methods by which positive epistasis has been previously estimated (i.e., competition of plasmid-carriers against plasmid-free cells[4]) may not measure the actual cost of plasmid coinfection. Compensatory evolution to ameliorate the cost of plasmids has been described in a wide range of plasmid systems and prevents plasmid loss by weakening negative selection against the plasmid backbone over time[17–20]. If the fitness costs of multiple plasmids have common mechanistic causes, it is possible that the same compensatory mechanisms could simultaneously ameliorate the costs of multiple plasmids which may in turn then promote stable plasmid coinfection[15, 19]. The role of positive selection for beneficial plasmid-encoded accessory gene functions in plasmid coinfection is less well studied. Positive selection could promote plasmid coinfection if the fitness benefits of carrying multiple plasmids outweighed the accumulative fitness costs, or, alternatively, may inhibit coinfection by selecting for consolidation of beneficial functions onto fewer replicons, accompanied by the loss of redundant, costly plasmid backbone(s)[13, 14]. Understanding the roles of these various mechanisms in plasmid coinfection requires experimental tests, but while the fitness costs of single- and co-infection by plasmids have been estimated, studies tracking the longer-term dynamics of plasmid coinfection in bacterial populations are lacking.

Here, we consider the experimental coinfection dynamics for two distantly-related, naturally co-occurring conjugative mercury-resistance plasmids—pQBR103 and pQBR57—originally isolated from a field-site in the UK[21, 22]. Each of the plasmids individually causes a substantial fitness cost in the host bacterium *Pseudomonas fluorescens* SBW25[23]. Both plasmids encode a near identical copy of the *mer* mercury-resistance operon encoded on a Tn5042 transposon [23, 24], which allows plasmid-bearers to reduce toxic Hg(II) to Hg, providing a fitness benefit to plasmid-carriers at increased Hg(II) concentrations[23]. Although SBW25(pQBR57) outcompetes SBW25(pQBR103) in the absence of mercury, this competitive hierarchy is reversed in mercury-containing environments[23]. This suggests that while pQBR57 imposes a lower fitness cost on SBW25, it also provides less of a fitness benefit in the presence of mercury, relative to pQBR103. Both plasmids are maintained in single-infected bacterial populations and, while this appears in each case to be linked to compensatory evolution [18, 25], unlike pQBR103, pQBR57 is capable of a high rate of conjugative transfer, which contributes to its survival and spread particularly in the absence of mercury[26, 27]. Multiple mechanisms of compensatory evolution have previously been described for the pQBR plasmids. Specifically, chromosomal compensatory mutations that occur either in *gacA/gacS* encoding a two-component global regulatory system or in *PFLU4242* encoding a hypothetical protein with two domains of unknown function, reduce the cost of these and other pQBR plasmids individually[17, 18].

Replicate laboratory populations of SBW25 that were originally either singly-infected or coinfected with the plasmids pQBR103 and pQBR57 were propagated by serial transfer with or without positive selection (i.e. addition of mercury (II) chloride) for approximately ~265 bacterial generations. We tracked bacterial population densities and the dynamics of mercury resistance over time and used multiplex-PCR to determine the plasmid-carriage status of mercury resistant clones. We show that plasmid coinfection was stable in populations without positive selection, whereas positive selection drove the loss of pQBR57 and the dominance of SBW25 carrying pQBR103-only. Loss of plasmid coinfection occurred despite compensatory evolution to ameliorate plasmid fitness costs and was caused by positive selection discriminating between the differential fitness benefits of the plasmids, retaining only the more beneficial plasmid.

## Material and Methods

### Bacterial Strains and Culture Conditions

Bacterial populations were grown in liquid Kings B (KB)[28] broth microcosms (6 ml of KB broth in a 30ml glass universal vial). These were incubated at 28° C and shaken at 180 rpm. To generate positive selection for mercury resistance, microcosms were supplemented with 40 μM Hg (II) chloride, as required. Bacterial colonies were obtained by plating serial dilutions onto KB agar. To select particular bacterial strains (described below), agar plates were supplemented with gentamycin (30 μg/ml), streptomycin (250 μg/ml), Hg (II) chloride (20 μM for plating and 100 μM for replica plating), kanamycin (25 μg/ml) or X-gal (75 μg/ml), as required.

Two isogenic *P. fluorescens* SBW25 strains[21] with chromosomal resistance markers [either gentamycin resistance (Gm^R^) or a streptomycin resistance with LacZ (Sm^R^*lacZ*)] were used to enable creation of transconjugants[23, 29, 30]. Derived *P. fluorescens* SBW25 deletion mutants for the *gacS* gene [SBW25-Gm^R^-ΔgacS; [18]] or the *PFLU4242* gene [SBW25-Gm^R^-ΔPFLU4242; [17]] were used to enable measurement of the effects of these genes on plasmid costs in competition experiments. An isogenic plasmid-free *P. fluorescens* SBW25 Gm^R^ strain with a chromosomal Tn5042 mercury resistance transposon derived from the pQBR103 plasmid [SBW25(Tn5042)] was used to enable measurement of the fitness cost of the plasmid backbones in competition experiments[27].

Two mercury resistance plasmids were used in this study that had been previously isolated from agricultural soil in Oxfordshire, UK: pQBR57 [21] and pQBR103 [22]. An isogenic variant of pQBR103 marked with an mCherry fluorescent protein gene and a kanamycin resistance gene (mCherryKm^R^) was used to enable selection of coinfected bacterial cells[31]. Plasmids were introduced to bacterial strains by conjugation using standard protocols[23, 32]. Briefly, transconjugants were selected by plating onto KB agar plates supplemented with 20 μM Hg (II) chloride or 25 μg/ml kanamycin and the relevant antibiotic [either gentamycin (30 μg/ml) or streptomycin (250 μg/ml)] as appropriate to select the recipient strain. Plasmid status of transconjugant colonies was determined by PCR, as previously described[18, 26].

### Selection experiment

Six replicate populations each of SBW25-Sm^R^*lacZ*(pQBR57+pQBR103-Km^R^), SBW25-Sm^R^*lacZ*(pQBR57), SBW25-Sm^R^*lacZ*(pQBR103-Km^R^) were propagated with or without positive selection (i.e., supplementation with 40 μM Hg (II) chloride or 0 μM Hg (II) chloride, respectively), and six replicate populations of the SBW25-Sm^R^*lacZ* plasmid-free control were propagated without Hg (II) chloride. (Note: plasmid-free populations cannot survive in 40 μM Hg (II) chloride.) Each replicate was founded by 60 μl of an overnight liquid culture initiated from a single independent colony previously streaked on KB agar. One percent of each population was serially transferred to fresh media every 48h for 40 transfers, resulting in approximately 265 bacterial generations. Every 10 transfers serial dilutions of each population were plated onto KB agar and incubated at 28° C to enumerate bacterial densities. These plates were replica plated onto KB agar supplemented with Hg (II) chloride 100 μM/ml to determine the frequency of mercury resistance (Hg^R^). Twenty-four mercury resistant colonies per population per time-point were chosen at random to determine the presence of each plasmid and Tn5042 by multiplex PCR. We used three set of primers to target the Mer-Tn5042 transposon [F-TGCAAGACACCCCCTATTGGAC, R-TTCGGCGACCAGCTTGATGAAC], the pQBR103-plasmid specific origin of replication *oriV* [F-TGCCTAATCGTGTGTAATGTC, R-ACTCTGGCCTGCAAGTTTC] and the pQBR57-plasmid specific *uvrD* gene [F-CTTCGAAGCACACCTGATG, R-TGAAGGTATTGGCTGAAAGG] [18, 26]. Briefly, a mixture of 1x GoTaq Green (Promega, WI USA) with 0.71 μM of each primer to detect pQBR103, 0.89 μM of each primer to detect pQBR57 and 0.36 μM of each primer to detect Tn5042 was used with the following thermocycle program: 95 °C 5’, 30 x (95 °C 30’, 58 °C 30’, 72 °C 1’), 72 °C 5’ [26].

### Competition experiments

Competition experiments were used to measure the fitness costs associated with carrying plasmid(s) against a range of competitor strains. In all cases, overnight cultures of competitors were mixed in a 1:1 ratio and diluted 100-fold into KB microcosms ± 40 μM Hg (II) chloride and incubated for 48h at 28°C with shaking at 180rpm. Starting and final densities of each marked strain were determined by plating onto KB agar supplemented with X-gal, and relative fitness (*w*) was calculated as previously described[18].

To measure the fitness cost of the plasmid backbones per se, that is once the fitness effect of the Tn5042 is accounted for, the plasmid-bearing strains [SBW25-Sm^R^*lacZ*(pQBR103-Km^R^) or SBW25-Sm^R^*lacZ*(pQBR57) or SBW25-Sm^R^*lacZ*(pQBR57+pQBR103-Km^R^)] were competed against SBW25-Gm^R^(Tn5042) both with and without Hg (II) chloride. Six replicates were performed per comparison.

To measure the fitness cost of plasmid coinfection we competed SBW25-Sm^R^*lacZ*(pQBR57+pQBR103-Km^R^) against either SBW25-Gm^R^(pQBR103-Km^R^) or SBW25-Gm^R^ (pQBR57) or SBW25-Gm^R^(pQBR57+pQBR103-Km^R^) as a control, both with and without Hg (II) chloride. Twelve replicates were performed per comparison. Normalised fitness was calculated by subtracting the mean of the control competition [i.e., SBW25-Sm^R^LacZ(pQBR57+pQBR103-Km^R^) versus SBW25-Gm^R^(pQBR57+pQBR103-Km^R^)].

To measure the fitness effect of putative compensatory mutations on plasmid carriage we competed SBW25-Gm^R^ or SBW25-Gm^R^-ΔgacS or SBW25-Gm^R^-ΔPFLU4242 carrying plasmid(s) (pQBR57 or pQBR103-Km^R^ or pQBR57+pQBR103-Km^R^) against plasmid-free SBW25-Sm^R^*lacZ*. Six replicates were performed per comparison.

### Genomic analysis

The whole genome sequence for at least one randomly chosen clone per population was obtained at the end of the selection experiment. For populations containing multiple mercury resistant genotype subpopulations—i.e., clones without mercury resistance or Tn5042-only or pQBR103-only or pQBR57-only or both pQBR57+pQBR103—and where these genotype(s) comprised at least 10% of the population, we obtained the whole genome sequence for one randomly chosen clone per subpopulation. Whole-genome sequencing was performed by MicrobesNG using a 250bp paired-end protocol on the Illumina HiSeq platform. Paired reads were aligned to the annotated ancestral genome sequence using Burrows-Wheeler Aligner[33] and duplicate reads were removed using picard (https://broadinstitute.github.io/picard/). Variants were called using GATK Haplotype Caller[34] and annotated using SnpEff[35]. Called variants were then filtered to remove low quality calls with either low coverage (< 12 reads per bp), low quality (scores <200) or low frequency of the alternative allele (< 90% of reads with alternative). In addition, as a complementary and confirmatory approach, variants were also called against the ancestral reference genome using the Breseq computational pipeline using the standard default settings [36]. All variants not called by both methods were validated visually using the alignment viewer IGV[37, 38]. All sequencing data are available on the Short Read Archive under accessions PRJEB38218 / ERP121615.

### Statistical analysis

The integral of mercury resistance frequency over time was calculated as the area under the curve using the AUC function of the ‘flux’ package in R and compared between treatments using Welch’s ANOVA due to unequal variances. Posthoc pairwise comparisons were performed using the Dunnetts T3 test. The integral of plasmid coinfection frequency over time was calculated as the area under the curve using the AUC function of the ‘flux’ package in R and compared between treatments using a Mann Whitney U test due to unequal variances. Relative fitness data from competition experiments was analysed using ANOVA and posthoc pairwise comparisons were performed using Tukey tests. Analyses were performed in R 3.6.1 [39] or Prism v8.1.2.

## Results

### Temporal dynamics of mercury resistance and plasmid carriage

To study the effect of positive selection on the dynamics of plasmid coinfection we propagated replicate populations of SBW25 carrying either both plasmids or pQBR103-alone or pQBR57-alone, both with and without selective levels of Hg(II) chloride by serial transfer for approximately 265 bacterial generations. We also propagated replicate plasmid-free control SBW25 populations without Hg(II) chloride. Mercury resistance (Hg^R^) was maintained near fixation under positive selection for all plasmid treatments, but without positive selection its frequency varied according to plasmid treatment (Figure S1; comparison of cumulative Hg^R^ frequency between plasmid treatments without Hg(II) selection: Welch’s ANOVA; W_2,6.685_ = 18.32, P = 0.0019). Without positive selection, whereas Hg^R^ remained at high frequency in the both-plasmids treatment, Hg^R^ declined in the single-plasmid treatments, and significantly so in the pQBR103-alone treatment (Dunnett’s T3 test for pairwise comparisons of plasmid treatments without Hg(II) selection: pQBR103:both P = 0.0044; pQBR57:both P = 0.4038; pQBR103:pQBR57 only P = 0.3157). The Hg^R^ phenotype indicates the presence of the Tn5042-encoded *mer* operon within the cell, which could be explained by the maintenance of one or both plasmids or by the relocation of the Tn5042 to the chromosome accompanied by plasmid loss[18, 26, 30]. We therefore used multiplex-PCR to determine the presence of the Tn5042 and of each plasmid. In the single-plasmid treatments, whereas plasmid-encoded Hg^R^ predominated in populations propagated without positive selection, we observed the invasion of plasmid-free cells carrying a chromosomal Tn5042 in some replicates with positive selection (Figure 1; Figure S2; 2 / 6 replicates of the pQBR57-alone treatment; 1 / 6 replicates of the pQBR103-alone treatment). Plasmid dynamics also varied with positive selection in the both-plasmids treatments. Plasmid coinfection was maintained at higher frequency in populations without positive selection compared to those propagated with positive selection (Figure 1; comparison of cumulative coinfection frequency: Mann Whitney U test, U = 0, P = 0.0022). This was driven by the loss of pQBR57 from initially coinfected cells under positive selection, such that plasmid-coinfected cells were replaced by cells carrying pQBR103-only in 4 out of 6 replicates (Figure S2). Taken together, these data suggest that positive selection inhibited plasmid coinfection and, furthermore, that the pQBR57-backbone was selected against more strongly than the pQBR103-backbone under positive selection for mercury resistance.

**Figure 1.**
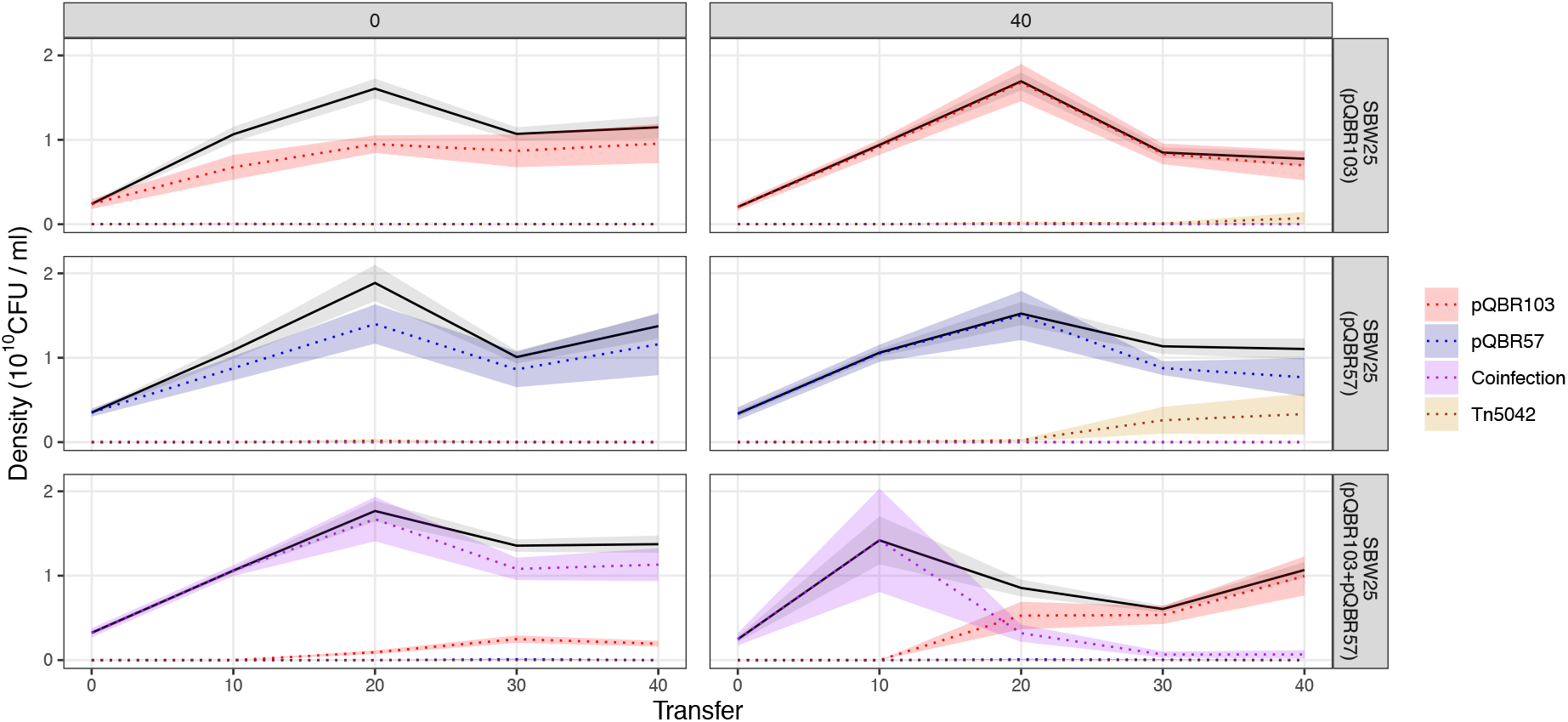
Population and mercury resistance genotype dynamics over time. Plots are facetted by treatment, horizontally by mercury treatment (0 μM or 40 μM Hg (II) chloride) and vertically by plasmid treatment (pQBR103-alone or pQBR57-alone or both pQBR103+pQBR57). Solid black lines show the mean (n = 6) ± standard error (grey shaded area) bacterial population density estimated from colony counts. Coloured dotted lines show the mean ± standard error (correspondingly coloured shaded area) population density of pQBR103-only carriers (red), pQBR57-only carriers (blue), coinfected pQBR103+pQBR57-carriers (purple), and chromosomal Tn5042 carrying plasmid-free cells (gold) inferred from their genotype frequencies. Plots for the population and genotype dynamics of individual replicates are shown in Figure S2. Raw data is provided in DataS1.

### Fitness effects of plasmid carriage

To understand the effect of positive selection on the relative fitness of the various Hg^R^ genotypes observed here—i.e., chromosomal Tn5042, either plasmid alone or both plasmids together—we performed a series of competition experiments. First, we tested how positive selection affected the cost of carrying the plasmid backbone(s) by competing SBW25 plasmid-bearers against an isogenic ancestral SBW25 encoding a chromosomal copy of the Tn5042 [SBW25(Tn5042)]. Here, in each pairwise competition both competitors are resistant to mercury, but the plasmid bearers must pay the additional fitness costs of maintaining the plasmid backbone(s). Plasmids bearers varied in fitness relative to SBW25(Tn5042) according to their plasmid complement: pQBR103 had a higher fitness cost than either pQBR57 or both plasmids together (Figure 2; ANOVA; plasmid main effect, F_2,28_ = 7.813 P = 0.002; Tukey pairwise contrasts: pQBR103:pQBR57 P = 0.0217; pQBR103:both P = 0.0019; pQBR57:both P = 0.6399). Moreover, the addition of Hg(II) chloride increased the fitness cost of plasmid carriage (mercury main effect, F_1,28_ = 31.298 P = 5.48 x 10^−6^), suggesting that once the benefit of mercury resistance is negated, Hg(II) chloride increased the costs of the plasmid backbones *per se*. Together, these data confirm previous studies reporting a higher cost of the pQBR103 backbone relative to the pQBR57 backbone[23] and, moreover, explain the loss of redundant plasmid replicons under positive selection seen here and in other studies[26, 31].

**Figure 2.**
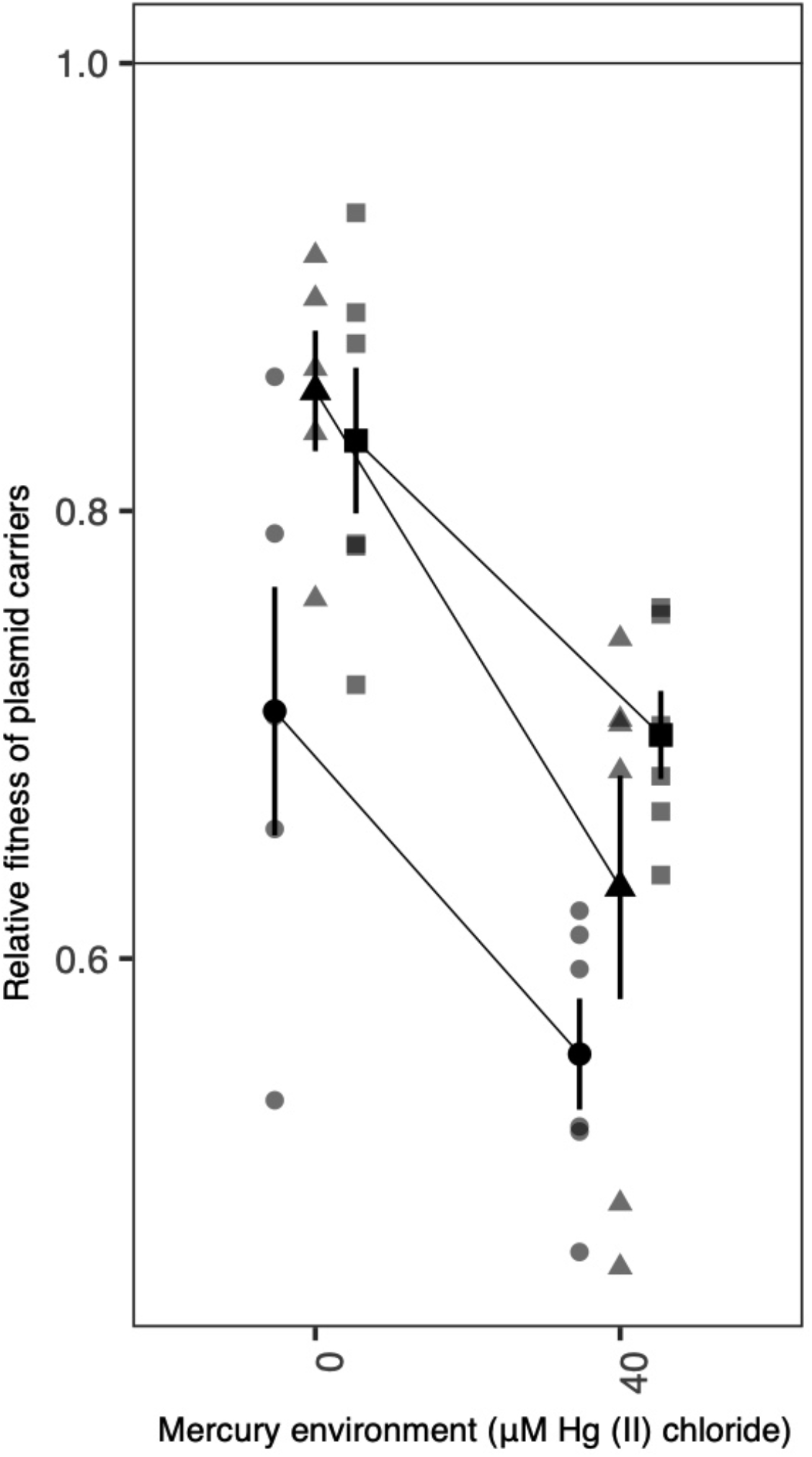
The fitness of plasmid carriers relative to SBW25(Tn5042) competed with or without Hg (II) chloride. Black symbols show mean ± standard error relative fitness of SBW25(pQBR103) (circles), SBW25(pQBR57) (triangles) or SBW25(pQBR103+pQBR57) (squares) against SBW25(Tn5042). Grey symbols in corresponding shapes show the individual replicate values (n = 5 or 6). Points are jittered to prevent over-plotting. All competition assays were six-fold replicated except for SBW25(pQBR103) versus SBW25(Tn5042) in 0 μM Hg (II) chloride and SBW25(pQBR57) versus SBW25(Tn5042) in 0 μM Hg (II) chloride, where one replicate each was lost due to contamination. Raw data is provided in DataS1.

Nevertheless, these data cannot explain the preferential loss of the pQBR57 backbone from coinfected cells that we observed. To further explore this, we next directly competed SBW25 coinfected with both plasmids against SBW25 carrying either of the plasmids alone, both with and without positive selection. We found that the relative fitness of coinfected SBW25 was lower when competed against SBW25(pQBR103) than against SBW25(pQBR57) (Figure 3; ANOVA; plasmid main effect, F_1,40_ = 7.438 P = 0.009435) and was reduced by the presence of Hg(II) in the media (mercury main effect, F_1,40_ = 5.727 P = 0.02149) while the interaction of these factors was marginally non-significant (plasmid:mercury interaction, F_1,40_ = 3.616 P = 0.06443). Post-hoc pairwise comparisons were consistent with positive selection favouring the loss of pQBR57 from coinfected cells as we observed in our serial transfer experiment: The addition of Hg(II) chloride further reduced the fitness of coinfected SBW25(pQBR57+pQBR103) when competed against SBW25(pQBR103) (Tukey pairwise comparison; with versus without Hg(II) chloride, P = 0.0203) but not when competed against SBW25(pQBR57) (Tukey pairwise comparison; with versus without Hg(II) chloride, P = 0.9967), such that the fitness of coinfected cells was higher against SBW25(pQBR57) than against SBW25(pQBR103) in the presence of Hg(II) chloride (Tukey pairwise comparison; P = 0.0113). Taken together and considering previous findings [23], these data suggest that although the pQBR103 backbone is costlier than the pQBR57 backbone, pQBR103 is more beneficial in mercury-containing environments than pQBR57. Thus, the preferential loss of pQBR57 from coinfected cells under positive selection is better explained by selection favouring the more beneficial plasmid rather than selection against the more costly plasmid backbone.

**Figure 3.**
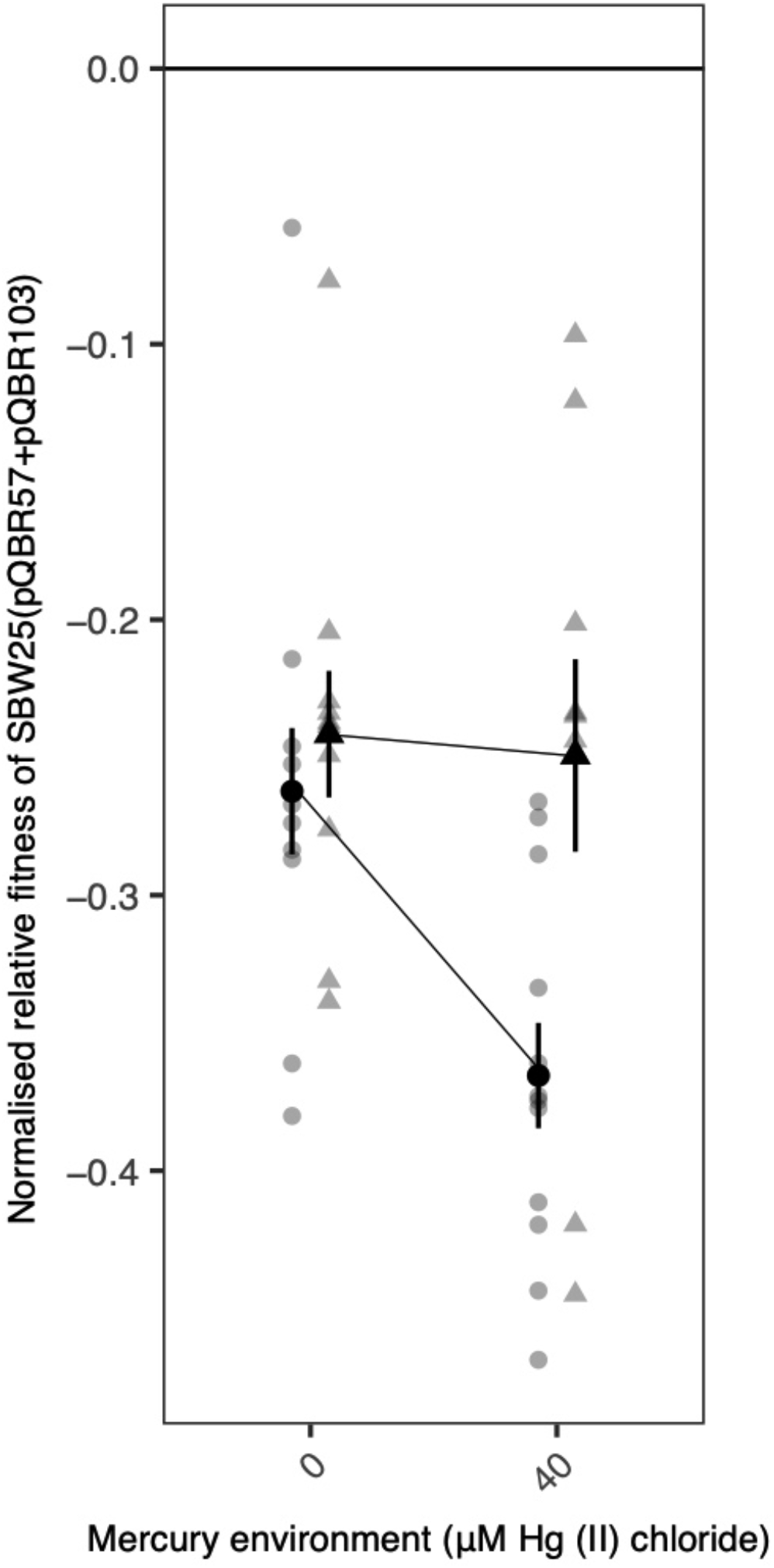
The fitness of coinfected plasmid carriers against singly-infected plasmid carriers competed with or without Hg (II) chloride. Black symbols show mean ± standard error normalised relative fitness of SBW25(pQBR103+pQBR57) against SBW25(pQBR103) (circles) or SBW25(pQBR57) (triangles). Grey symbols in corresponding shapes show the individual replicate values (n = 10 or 12). Points are jittered to prevent over-plotting. Competition assays against SBW25(pQBR103) were twelve-fold replicated, whereas competition assays against SBW25(pQBR57) were ten-fold replicated due to the loss of replicates to contamination. Raw data is provided in DataS1.

### Compensatory mutations and their fitness effects

A potential alternative explanation for the contrasting patterns of plasmid maintenance among treatments would be if there existed differences in the propensity for compensatory evolution to occur according to plasmid content or positive selection. To test this, we obtained the whole genome sequences of one randomly chosen evolved clone per Hg^R^ genotype present at greater than 10% frequency per population at the end of the serial transfer experiment. There was no evidence for differential compensatory evolution according to treatment: all sequenced plasmid-bearing evolved clones carried a mutation in a known compensatory locus. Specifically, we observed mutations in *gacA, gacS* or *PFLU4242*, or in regions immediately upstream of compensatory loci (Figure 4; data for all sequenced clones provided in Figure S3-S9). By contrast, mutations at these loci were never observed in evolved clones from the plasmid-free control populations. To confirm that loss of either *gacS* or *PFLU4242* was sufficient to ameliorate the cost of plasmid co-infection, we compared the effect of compensatory mutations on the fitness of plasmid bearers carrying either one or both plasmids relative to the plasmid-free ancestor using competition experiments. Consistent with a role for these genes in compensatory evolution, deletion of either *gacS* or *PFLU4242* reduced the fitness cost of carrying plasmids (Figure 5). However, while deletion of either gene completely ameliorated the cost of carrying either both plasmids or pQBR103 alone, only deletion of *PFLU4242* completely ameliorated the cost of carrying pQBR57 alone, whereas deletion of *gacS* offered only partial amelioration (ANOVA; strain by plasmid interaction, F_4, 39_ = 2.628, P = 0.049). Thus, compensatory evolution occurred in all treatments and was effective at ameliorating the additional cost of carrying multiple plasmids. While this may explain the stability of plasmid coinfection without positive selection, it cannot explain the decline in coinfection due to loss of pQBR57 with positive selection, which instead appears to have been driven by the differential benefits of the plasmids in the presence of Hg(II) chloride.

**Figure 4.**
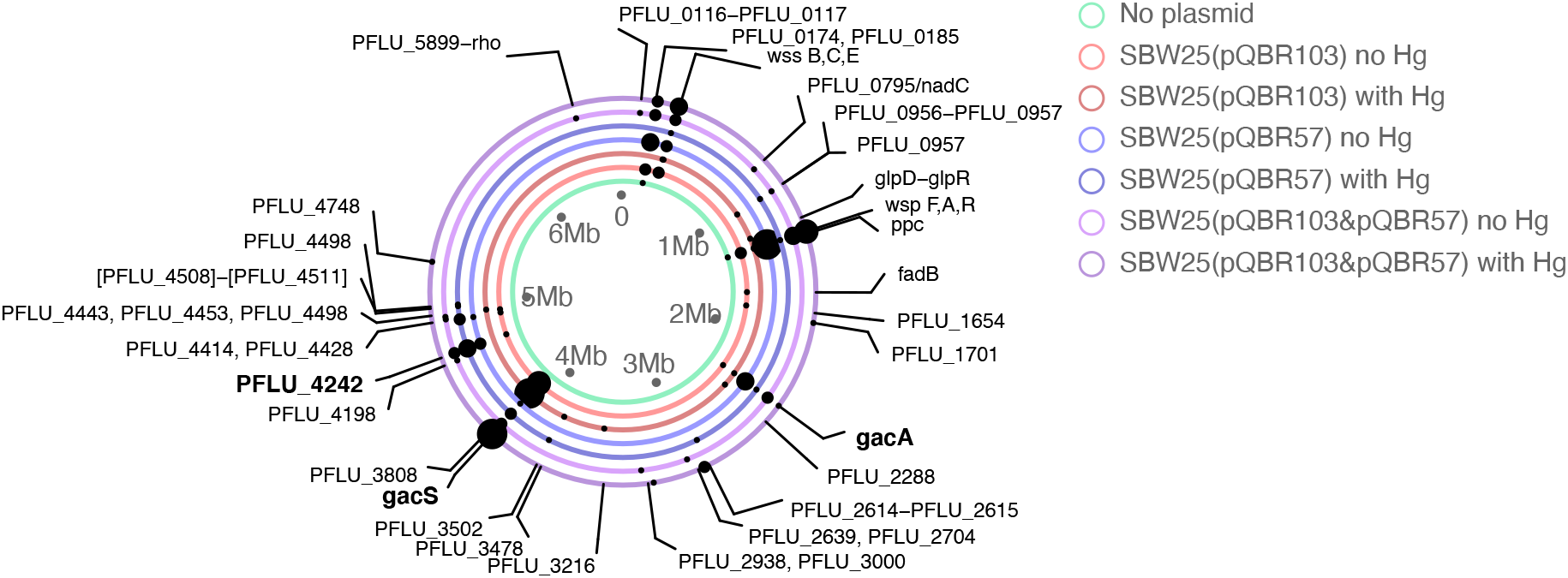
Parallel evolution plot summarising the mutations arising in evolved clones per treatment. Rings represent the SBW25 chromosome plotted separately for each treatment as indicated by the colour of the ring (see legend). Dots indicate genetic loci where mutations (excluding hypermutator clones) were observed in that treatment; the size of the dot corresponds to the number of replicate populations where a mutation at that locus occurred per treatment. Loci previously associated with compensatory mutations are highlighted in bold. In populations where more than one plasmid-bearing clone was sequenced, only mutations present in the dominant genotype are shown here. Hyper-mutator clones are not shown to avoid over-plotting. Plots for all sequenced clones are provided in Figure S3. The full table of called sequence variants is provided in DataS2.

**Figure 5.**
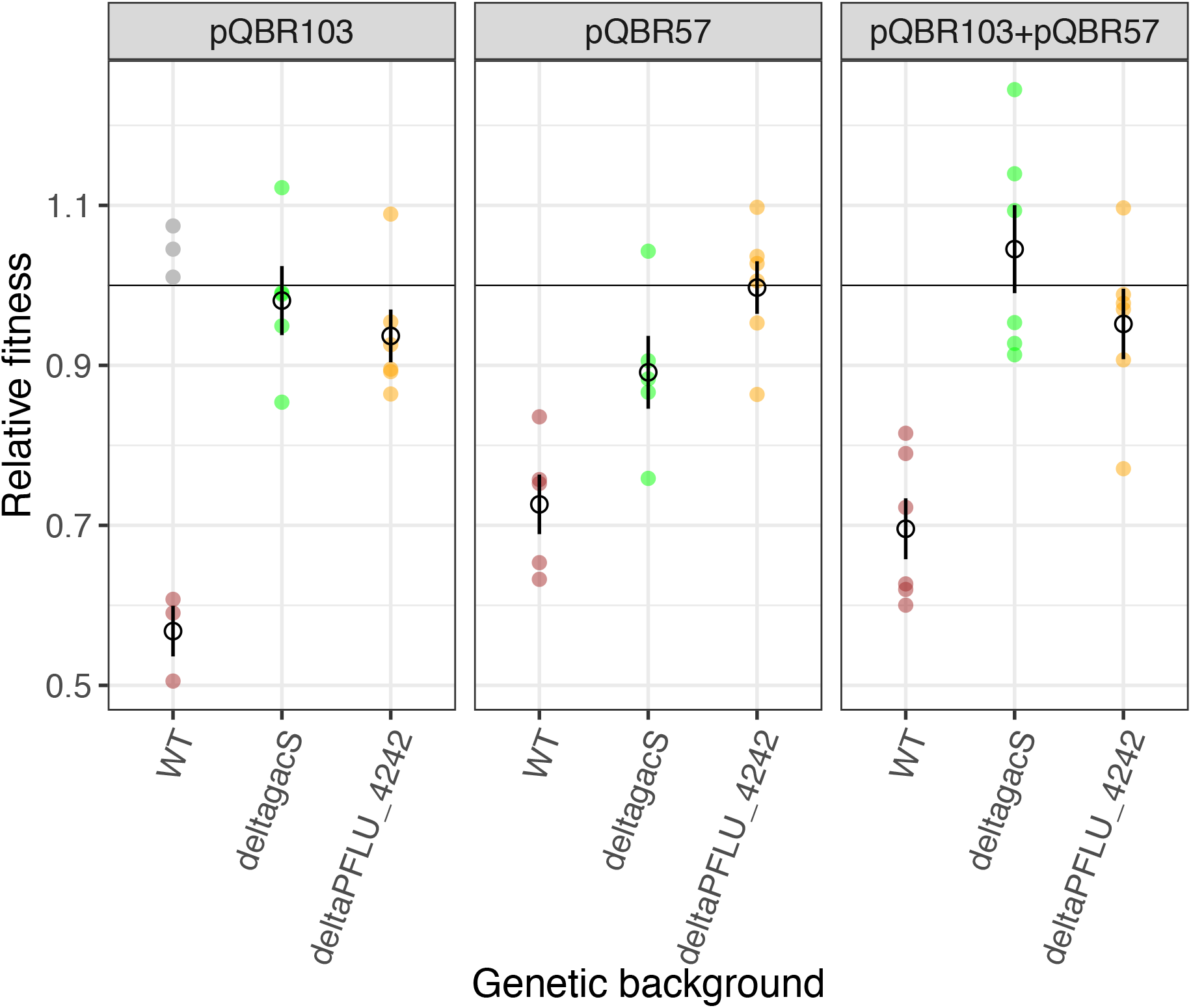
Compensatory mutations ameliorate the fitness costs of plasmid carriage. Plots are facetted horizontally by the plasmid status of the plasmid-carrying competitor. Open symbols show the mean (n = 6) ± standard error of fitness for plasmid carrying genotypes relative to plasmid-free SBW25. Coloured filled dots denote individual replicate values for SBW25 (maroon), SBW25-ΔgacS (green) or SBW25-ΔPFLU4242 (orange) genotypes. Outliers (grey) were excluded from the statistical analysis. Raw data is provided in DataS1.

## Discussion

Plasmid coinfection is more commonly observed in bacterial genomes than is expected by chance[4] which is surprising given the high fitness costs usually associated with plasmid acquisition[1, 2, 11, 13]. Here, we show that coinfection by plasmids encoding the same functional trait, namely mercury resistance, was inhibited by positive selection for this trait. With positive selection, coinfection declined due to loss of pQBR57 from originally coinfected cells, whereas without positive selection, plasmid coinfection was stably maintained. This difference could not be explained by the fitness costs of plasmid carriage, firstly because coinfection was maintained in environments where the net cost of plasmid carriage was highest (i.e. without Hg (II) chloride), and secondly because compensatory mutations to ameliorate these costs occurred across all treatments. Instead, the loss of pQBR57 from originally coinfected cells under positive selection could be explained by the differential fitness benefits provided by the two plasmids in the presence of Hg (II) chloride. Addition of Hg (II) chloride reduced the fitness of coinfected cells when competing against pQBR103-carriers but not when competing against pQBR57-carriers. Consistent with previous data from singly-infected cells[23], we show that this occurs because although the pQBR103 backbone is costlier, it provides a greater benefit in the presence of Hg (II) chloride. Thus, where multiple plasmids encode the same functional trait, under positive selection this redundancy can drive loss of the less beneficial plasmid.

Loss of a redundant plasmid replicon by coinfected cells under positive selection is conceptually similar to the replacement of single-infected plasmid bearers by the chromosomal Tn5042, which was also only observed under positive selection. In both cases, redundancy allows for loss of one of the replicons encoding resistance. It may be the case that if our experiment was run for even longer, the initially coinfected populations would also eventually consist only of cells with a chromosomal Tn5042. Indeed, our relative fitness data show that SBW25(Tn5042) outcompeted both single- and double-infected plasmid bearers in mercury-containing environments. This outcome may have been delayed (or prevented) because the loss of both plasmids by segregation is likely to occur with low probability and/or because the fitness difference between chromosomal and plasmid-encoded mercury resistance was reduced by compensatory evolution. The key difference between environments with versus without positive selection, is that, in the former, both the benefit and the cost of the plasmids contribute to their fitness effect, whereas in the latter only their costs do. This appears to increase the fitness difference between coinfected cells and those carrying only the more beneficial plasmid. Selection for plasmid benefits may therefore be more efficient than negative selection against plasmid costs, which becomes less efficient over time due to compensatory evolution to ameliorate the plasmid fitness cost.

Previous studies suggest that the positive epistasis of plasmid fitness costs enables plasmid coinfection[4]. Our data provide contradictory evidence both for and against this idea. When competing against a plasmid-free genotype [SBW25(Tn5042)], SBW25 coinfected with both plasmids showed a lower fitness cost than expected from the individual fitness costs of each plasmid alone. This is consistent with positive epistasis. However, when coinfected SBW25 was directly competed against SBW25 singly-infected with either of the plasmids, we measured an appreciable additional cost of plasmid coinfection. This is not consistent with positive epistasis. It seems likely that direct competition of coinfected versus singly-infected cells offers the most accurate method to measure the fitness effect of carrying a second plasmid replicon. Whereas, competing singly-infected and coinfected cells separately against a plasmid-free competitor seems likely to underestimate the actual cost of carrying a second plasmid. Importantly, the study by San Millan and colleagues (2014) used a set-up similar to our first set of experiments, namely separate competition against a common plasmid-free strain[4], a method that our data suggest is likely to underestimate the actual fitness cost of coinfection. Our data when taken together do not, therefore, support a role for positive epistasis in explaining plasmid coinfection and urge caution when interpreting epistasis from relative fitness data obtained against plasmid-free competitors.

At present, it is unclear why the two plasmids differ in the benefit they provide to SBW25 under mercury positive selection. Both plasmids possess one copy of the Tn5042 transposon encoding the Mer operon, which provides mercury resistance. These Tn5042 sequences are identical except for a single base pair difference in *merR*, a repressor controlling expression of the Mer operon[23]. Low concentrations of Hg(II) are bound by MerR, causing a conformational change of the protein which relieves repression, allowing expression of the Mer operon, which imports Hg(II) into the cell and reduces it to Hg[40, 41]. It is possible that the single base pair difference in the *merR* sequence between the pQBR103 and pQBR57 copies of the Tn5042 could change the sensitivity of the MerR repressor leading to altered expression of the Mer operon. For example, by detoxifying the environment through reduction of Hg(II) to Hg, mercury resistance benefits all neighbouring cells [42], thus a reduced affinity for Hg(II) could accrue fitness benefits when in competition with strains encoding MerR with a higher affinity and thus earlier expression of the Mer operon. Alternatively, the plasmids differ extensively in their gene content, and it is possible that epistatic interactions between these variable genes and the Mer operon could underlie the observed differential fitness benefits. Resolving this is beyond the scope of this paper but identifying the causes of differences in plasmid benefit will be a focus of future work.

We observed mutations in three loci previously associated with compensatory evolution for pQBR plasmids. Specifically, mutations to either of the genes encoding the GacAS two-component global regulatory system[17, 18], or the gene *PFLU4242* of unknown function[17], were sufficient to ameliorate the fitness costs associated with coinfection by both pQBR103 and pQBR57. Similarly, deletion of either locus could completely ameliorate the cost of pQBR103, whereas amelioration of pQBR57 was more complete via deletion of *PFLU4242* than *gacS*. Together this suggests that these plasmids, despite being genetically divergent, cause their associated fitness costs through a similar (or a shared) mechanism. Shared targets of compensatory evolution among divergent plasmids have been reported in other plasmid-host systems[15, 19], for example compensatory mutations in chromosomal helicases[19]. Consistent with previous work on other host-plasmid systems[15, 19], our findings suggest that compensatory evolution could indeed promote plasmid coinfection by reducing the cost of multiple plasmids simultaneously. This has the effect of reducing the efficiency of negative selection against plasmid costs over time. It is notable, however, that although compensatory evolution ameliorating the fitness costs of plasmid coinfection occurred in both the mercury-containing and mercury-free environments, plasmid coinfection was only stable in the mercury-free environment (at least for the duration of this study). This occurs because positive selection remains efficient even after compensatory evolution occurs, allowing discrimination of differential fitness benefits among functionally redundant plasmid replicons.

These data reveal that coinfecting plasmids are unlikely to be maintained under positive selection if they encode redundant functions and there are differences in the fitness benefits these accessory genes provide. Moreover, while compensatory evolution can promote plasmid coinfection in the absence of positive selection, it does not do so in the presence of positive selection where not only the cost but also the benefit of the plasmid is subject to selection. These findings suggest therefore that widespread plasmid coinfection is likely to be explained either by positive selection for accessory gene functions being rare in nature, or by coinfecting plasmids encoding non-redundant accessory gene functions. These findings have implications for understanding the forces structuring bacterial genomes[1, 5–7], and suggest a process whereby recurrent phases of genome expansion and contraction are driven by variable positive selection: Multiple redundant replicons can be acquired, and their costs ameliorated by compensatory evolution thus allowing genome expansion between bouts of periodic positive selection, which, when they subsequently occur, then select reduced genomes containing only the highest benefit replicon(s).

## Supporting information

DataS1

DataS2

## Acknowledgements

This work was funded by grants from the European Research Council (ERC Starting Grant agreement 311490), the Natural Environment Research Council (NE/R008825/1), the Biotechnology and Biological Sciences Research Council (BB/R006253/1), and the Leverhulme Trust (PLP-2014-242) to MAB. EH is funded by an Independent Research Fellowship from the Natural Environment Research Council. JPJH is funded by a Tenure Track Fellowship from the University of Liverpool.

